# Mu-opioid receptor knockout on Foxp2-expressing neurons reduces aversion-resistant alcohol drinking

**DOI:** 10.1101/2023.11.29.569252

**Authors:** Harrison M. Carvour, Charlotte A. E. G. Roemer, D’Erick P. Underwood, Edith S. Padilla, Oscar Sandoval, Megan Robertson, Mallory Miller, Natella Parsadanyan, Thomas W. Perry, Anna K. Radke

## Abstract

Mu-opioid receptors (MORs) in the amygdala and striatum are important in addictive and rewarding behaviors. The transcription factor *Foxp2* is a genetic marker of intercalated (ITC) cells in the amygdala and a subset of striatal medium spiny neurons (MSNs), both of which express MORs in wild-type mice and are neuronal subpopulations of potential relevance to alcohol-drinking behaviors. For the current series of studies, we characterized the behavior of mice with genetic deletion of the MOR gene *Oprm1* in *Foxp2*-expressing neurons (Foxp2-Cre/Oprm1^fl/fl^). Male and female Foxp2-Cre/Oprm1^fl/fl^ mice were generated and heterozygous Cre+ (knockout) and homozygous Cre-(control) animals were tested for aversion-resistant alcohol consumption using an intermittent access (IA) task, operant responding for a sucrose reward, conditioned place aversion (CPA) to morphine withdrawal, and locomotor sensitization to morphine. The results demonstrate that deletion of MOR on *Foxp2*-expressing neurons renders mice more sensitive to quinine-adulterated ethanol (EtOH). Mice with the deletion (vs. Cre-controls) also consumed less alcohol during the final sessions of the IA task, responded less for sucrose under an FR3 schedule, and were less active at baseline and following morphine injection. *Foxp2*-MOR deletion did not impair the ability to learn to respond for reward or develop a conditioned aversion to morphine withdrawal. Together, these investigations demonstrate that *Foxp2*-expressing neurons may be involved in escalation of alcohol consumption and the development of compulsive-like alcohol drinking.

## 1. Introduction

The endogenous opioid system is an important regulator of rewarding experiences, including pleasure and relief of pain. As such, endogenous opioid peptides and their receptors are implicated in addictive behaviors across a variety of drug classes (Gianoulakis, 2009). The mu-opioid receptor (MOR), which is coded for by the *Oprm1* gene, is widely expressed throughout the central nervous system and directs inhibitory mechanisms both pre- and postsynaptically (Erbs et al., 2015). MORs are strongly implicated in alcohol drinking behaviors, including consumption, preference, and the acute stimulant and anxiolytic actions of ethanol (EtOH) (Becker et al., 2002; Ghozland et al., 2005; Hall et al., 2001; Heyser et al., 1999; Méndez and Morales-Mulia, 2008; Ray et al., 2013). Specifically, pharmacological blockade of MORs has been shown to reduce alcohol-seeking behaviors in mice (Gilpin et al., 2008; Heyser et al., 1999; Hyytiä and Kiianmaa, 2001; Marinelli et al., 2010). However, it has not yet been determined how specific subpopulations of MORs contribute to various alcohol-drinking behaviors.

The amygdala and striatum are two brain regions that have been identified as critical sites for the influence of MOR activation on addictive behaviors (Ben Hamida et al., 2019; Foster et al., 2003; Lam et al., 2008; Marinelli et al., 2010; Olive et al., 2001; Perry and McNally, 2013; Richard and Fields, 2016; Zhang and Kelley, 2002). Within each of these nuclei, MORs are expressed on several subpopulations of neurons. In the amygdala, MOR expression is densest in the *Foxp2*-expressing intercalated (ITC) cells (Campbell et al., 2009; Likhtik et al., 2008). ITC cells are GABAergic interneurons that are critical regulators of extinction learning and increase GABA release when exposed to ethanol (EtOH) (Hagihara et al., 2021; Likhtik et al., 2008; Silberman et al., 2009). In the striatum, MORs are found in medium spiny neurons (MSNs), which also express the transcription factor *Foxp2*. Approximately 65-75% of *Foxp2*-expressing MSNs co-express dopamine type 1 receptors (D1Rs) and 20-25% co-express dopamine type 2 receptors (D2Rs) (Fong et al., 2018). Both D1R and D2R-expressing neurons are thought to be involved in rewarding and addictive behaviors, with D1R-expressing neurons being involved in approach behaviors and D2R-expressing neurons being more involved in aversion and punishment (Hikida et al., 2016; Lobo and Nestler, 2011). Thus, *Foxp2*-MORs could be a subpopulation of relevance to alcohol-drinking behaviors.

Aversion-resistant, or “compulsive,” alcohol drinking in rodents is frequently used to model the tendency for individuals with Alcohol Use Disorder (AUD) to drink despite negative consequences. To test aversion resistance, alcohol is paired with an aversive outcome such as a footshock or the bitter tastant quinine. Aversion-resistant alcohol drinking recruits cortical and striatal circuits (Arnold et al., 2023; Chen and Lasek, 2020; Halladay et al., 2020; Radke et al., 2017; Seif et al., 2013; Siciliano et al., 2019), including the nucleus accumbens (NAc) core (Seif et al., 2013; Sneddon et al., 2021). In a previous study, we demonstrated that non-specific chemogenetic inhibition of NAc core neurons reduced aversion-resistant alcohol consumption but that targeted inhibition of D1- or D2-receptor-expressing medium spiny neurons (MSNs) did not (Sneddon et al., 2021). These findings suggest that aversion-resistant behaviors require a subpopulation of striatal neurons not captured by the traditional D1 vs D2 dichotomy. NAc *Foxp2*-expressing neurons have been shown to contribute to appetitive and aversive learning (He et al., 2024) and, as noted above, express both D1R and D2R. As such, these neurons are a candidate for regulating aversion-resistant drinking.

We set out to characterize the contributions of *Foxp2*-MORs to alcohol consumption, preference, and aversion-resistance using a transgenic line of mice with genetic deletion of the *Oprm1* gene in *Foxp2*-expressing neurons. To further characterize the line and the potential contribution of *Foxp2*-MORs to other behavioral manifestations of reward and aversion, we also examined operant responding for a sucrose reward, morphine withdrawal-induced aversion, and morphine-induced locomotion. Our results suggest that MOR deletion on *Foxp2*-expressing neurons reduces aversion-resistant consumption of EtOH.

## 2. Materials and Methods

### 2.1. Subjects

Male and female mice with genetic deletion of *Oprm1* in *Foxp2* neurons were generated by crossing B6.129-Oprm1tm1.1 Cgrf/KffJ by B6.C (strain #030074) and Foxp2tm1.1(cre)Rpa/J lines of mice (strain #030541) (Jackson Laboratory, Bar Harbor, ME) (**Figure 1A**). Following three crosses, the Foxp2-Cre/Oprm1^fl/fl^ line was established and Cre-females were bred with Cre+ males to produce offspring that were Cre- or Cre+. For all experiments, groups were balanced across sex and genotype. Mice were housed at Miami University in standard shoe box rectangular mouse cages (18.4 cm x 29.2 cm x 12.7 cm) with food (LabDiet 5001) and reverse-osmosis (RO) water *ad libitum*. For drinking experiments, cage lids were designed to allow for two test bottles to be present at the same time, with food placed in the cage lid troughs. The animals resided in a temperature-controlled room with a 12:12 hour light/dark cycle. All procedures were conducted in accordance with the Guide for the Care and Use of Laboratory Animals and were approved by the Miami University Animal Care and Use Committee.

**Figure 1.**
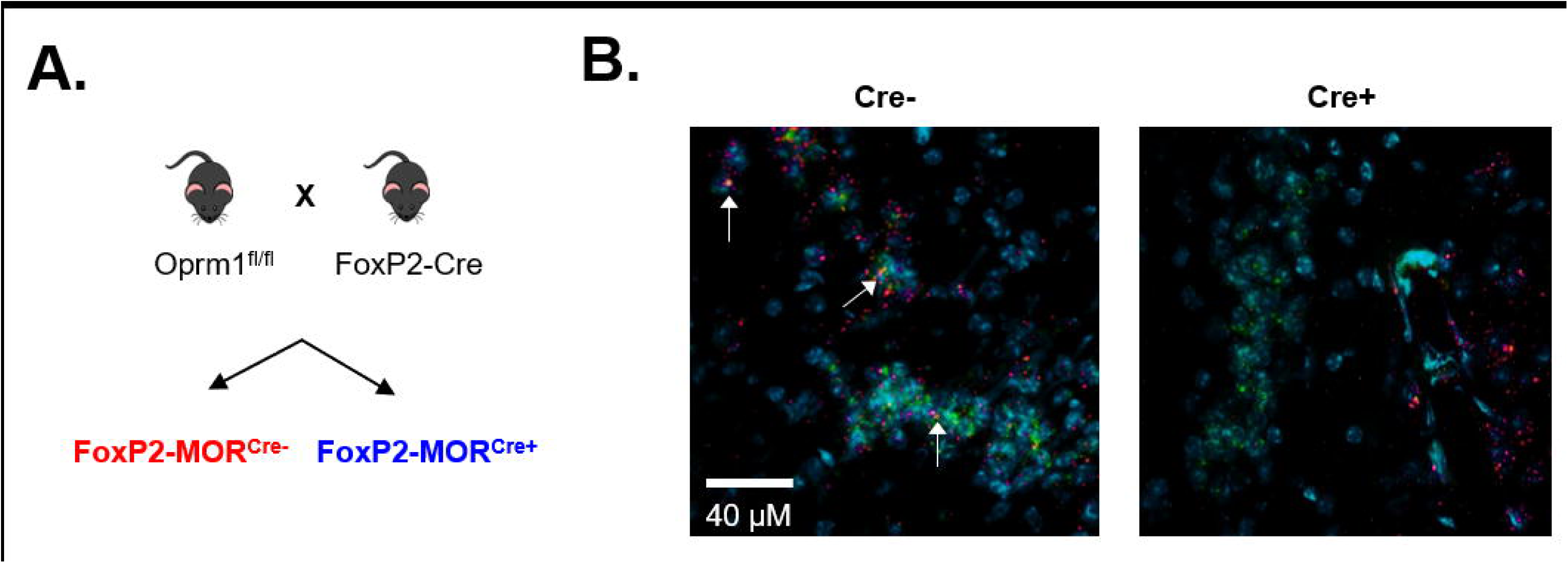
Selective genetic deletion of the mu-opioid receptor *Oprm1* gene on *Foxp2*-expressing neurons. **A)** Heterozygous Foxp2-Cre males (Foxp2tm1.1(cre)Rpa/J, strain #030541) were bred with homozygous *Oprm1* floxed females (B6.129-Oprm1tm1.1 Cgrf/KffJ by B6.C, strain #030074) to produce a Cre-mediated deletion of the MOR gene (*Oprm1*) on *Foxp2*-expressing neurons. Foxp2-Cre/Oprm1^fl/fl^ Cre-offspring had intact MORs while Cre+ offspring were missing MORs on *Foxp2*-expressing neurons. **B)** Fluorescent *in situ* hybridization confirmed *Oprm1* deletion in *Foxp2* neurons in the amygdala. Green = *Foxp2*, magenta= *Oprm1*, cyan= DAPI nuclear stain.

### 2.2. Drugs

EtOH (20%) was made volume/volume in RO water. Sucrose solutions (10%, 5%, 2%) were made weight/volume in RO water. Quinine hemisulfate (Q1250-50G, Millipore-Sigma, St. Louis, MO) was prepared weight/volume in 20% EtOH, RO water, or sucrose. For intermittent access, solutions were made fresh every day. For operant conditioning, sucrose solutions were prepared every three days.

Morphine sulfate was purchased from Henry Schein (Melville, NY, USA, #1099591) as 50 mg/ml vials. Injectable solutions of 1 mg/mL were prepared by diluting stock solution in sterile 0.9% saline solution (Henry Schein, Melville, NY, USA, #1048688). Naloxone hydrochloride (Sigma-Aldrich, St. Louis, MO, USA, #N7758) was dissolved in sterile saline to a concentration of 0.25 mg/mL. Prepared solutions were kept in a 4℃ refrigerator. Solutions were injected i.p. at a volume of 10 mL/kg.

### 2.3. Fluorescent in situ hybridization

Validation of *Oprm1* deletion in the Foxp2-Cre/Oprm1^fl/fl^ line (n = 8 Cre- and 8 Cre+) was conducted using RNAscope (Advanced Cell Diagnostics, Newark, CA, USA). Brains from adult mice were extracted and flash frozen. A cryostat was used to collect sections (15 μM) containing the amygdala. Sections were mounted on slides and fixed in chilled formalin and processed using probes for Mm-*Foxp2* (channel 1, cat#-C1) 428791and Mm-*Oprm1* genes (channel 2, cat#315841-C2). Slides were counterstained with DAPI and visualized using a Leica AX70 microscope (**Figure 1B**).

### 2.4. Intermittent access alcohol drinking

48 Foxp2-Cre/Oprm1^fl/fl^ mice were presented with 20% EtOH in a 24-hour two-bottle choice intermittent access (IA) paradigm. Mice were separated and individually housed three days prior to the start of the experiment. Two empty control cages were also used to account for fluid loss due to evaporation or spillage. Mice received a bottle of 20% EtOH and a bottle of RO water on Mondays, Wednesdays, and Fridays. For EtOH and RO water sessions, mice were weighed before bottles were presented. On Tuesdays, Thursdays, Saturdays, and Sundays mice received two bottles of RO water only. Bottles were weighed 24 h later for all sessions, except Saturday and Sunday. On Mondays, bottles were also weighed 30 min after presentation to assess binge-like consumption.

After 12 sessions (4 weeks) of EtOH drinking, quinine, a bitter tastant, was added to the EtOH bottles in escalating concentrations (10, 100, 200 mg/L) to test aversion-resistant drinking. Following the quinine sessions, mice received RO water for one week before beginning a test of quinine sensitivity. For this test, mice had a choice of water and water with quinine in escalating concentrations (0, 1, 10, and 100 mg/L). Data from only 32 animals are included for quinine sessions, as 16 animals mistakenly received the wrong quinine concentrations during testing.

### 2.5. Operant responding for sucrose

To examine behavioral responses for a non-drug solution and learning, 32 Foxp2-Cre/Oprm1^fl/fl^ mice were tested for sucrose responding using an operant conditioning paradigm in a standard mouse conditioning chamber (Med Associated, ENV-307A, Fairfax VT, USA) (as described in (Sneddon et al., 2020)). Nose poke data were recorded using Med-PC V Behavioral Control Software (Med Associates Inc., Fairfax, VT, USA). Prior to response training, mice were restricted to 85% of their free-feeding body weight to promote engagement with the task. Body weights were recorded before each operant session and food was given immediately after the session ended.

Upon reaching their target body weight, mice began response training, in which active nose poke responses resulted in delivery of 50 µl of a 10% sucrose solution. Responses on the inactive nose poke hole were inconsequential. Rewards were delivered on a Fixed Ratio 1 (FR1) schedule during a 30 min session. After 10 training sessions, mice underwent a sucrose fading procedure where they were given 10%, 5%, and 2% sucrose for 3 sessions each. Next, aversion-resistant responding was tested using 2% sucrose. Mice were trained to respond on an FR3 schedule for 3 additional sessions, to encourage an increased response rate for 2% sucrose. Escalating concentrations of quinine (1, 10, 100, and 200 mg/L) were added to the 2% sucrose solution over 4 sessions, 1 session for each concentration.

### 2.6. Conditioned place aversion

To determine whether Foxp2-MOR deletion enhanced other forms of aversion, 14 Foxp2-Cre/Oprm1^fl/fl^ mice were tested for morphine withdrawal-induced conditioned place aversion (CPA). The place conditioning apparatus (Panlab, Barcelona, Spain) was made up of three chambers distinguished by different designs on the wall and different floor textures, with the middle chamber being neutral and not used for conditioning. A camera mounted above the apparatus was used to record the test sessions. All sessions were 25 min in duration and were separated by 24 h.

The experiment began with a pre-conditioning test during which mice freely explored both sides of the apparatus. Following the pre-conditioning test, mice underwent four conditioning sessions. For two sessions, mice received an injection of morphine (10 mg/kg) followed 30-min later by an injection of naloxone (2.5 mg/kg). Mice were confined to the withdrawal-paired conditioning chamber immediately after naloxone injection. For the other two sessions, mice received two injections of saline separated by 30 min before being confined to the other conditioning chamber. The conditioning chamber paired with withdrawal and the order of the conditioning sessions was counterbalanced across animals. After all conditioning sessions were completed, mice freely explored the apparatus for 25 min in a post-conditioning test.

### 2.7. Morphine-induced locomotion

To assess the effects of *Foxp2*-MOR deletion on baseline and morphine-induced locomotion, 23 Foxp2-Cre/Oprm1^fl/fl^ mice were tested in activity chambers (14 in x 14 in x 8 in) that tracked movement via beam breaks (Omnitech Electronics, Inc, Columbus, OH). One week prior to the experiment, mice were given saline injections to habituate them to handling. Mice were tested for locomotion over two sessions separated by 24 h. The first session was 60 min in duration. For the second session, mice received one injection of 20 mg/kg of morphine 30 min prior to entering the locomotor chamber and activity was monitored for 120 min.

### 2.8. Data analysis

For intermittent access, bottle weights were expressed as g of EtOH consumed per kg of body weight. The amount of solution consumed was measured by calculating *=(Initial Bottle Weight - Final Bottle Weight) - Average Dummy Bottle Weight*. Preference was measured by calculating *=(Volume of solution/Volume of solution + Water consumption)*100*. For quinine sessions, baseline EtOH consumption was determined by averaging the last three sessions before adding quinine (sessions 10, 11, and 12). For the quinine sensitivity test, consumption at 0 mg/L was averaged across both bottles to establish a baseline. This was done because some mice were observed to drink exclusively from one bottle on the 0 mg/L session. For operant experiments, responses at the active and inactive nose-poke holes were averaged across the three sessions for each sucrose solution delivered (10%, 5%, and 2%).

Place conditioning data were expressed as time spent (s) and distance traveled (cm) on the withdrawal-paired side of the apparatus during the pre- and post-conditioning tests. For the locomotor tests, total distance traveled (cm) was measured.

For all experiments, data were first analyzed with sex as a variable but later collapsed for presentation and further analysis as sex did not interact with genotype on any behavioral measure. Data were analyzed using two-way repeated measures (RM) ANOVA with genotype as the between-subjects variable and session, concentration, or time as the within-subjects variable. For cases where sphericity was violated (ε < 0.75), the Greenhouse-Geisser correction was used. In instances where values were missing due to spillage, a Mixed Effects ANOVA was used. For operant responding, a three-way RM ANOVA was used to assess responding on the active vs. inactive nose-poke hole. Aversion resistance was defined *a priori* as the absence of a significant change from baseline (= 0 mg/L) and was tested using a Dunnett’s test. All data were expressed as mean ± standard error of the mean and all analyses were performed using GraphPad Prism (v 10) software (La Jolla, CA, USA).

## 3. Results

### 3.1. Intermittent access alcohol drinking

*Foxp2*-MOR deletion did not affect EtOH consumption or preference at 24 h (**Figure 2A-B**). For consumption, RM two-way ANOVA revealed a significant main effect of session [F_(4.981, 229.1)_ = 2.492, p < 0.05], but no significant main effect of genotype [F_(1, 46)_ = 1.510, p > 0.05] or session x genotype interaction [F_(11, 506)_ = 1.043, p > 0.05]. Similarly, for EtOH preference, the main effect of session was significant [F_(6.443, 296.4)_ = 2.544, p < 0.05], but the main effect of genotype [F_(1, 46)_ = 0.1627, p > 0.05] and session x genotype interaction [F_(11, 506)_ = 0.6708, p > 0.05] were not significant.

**Figure 2.**
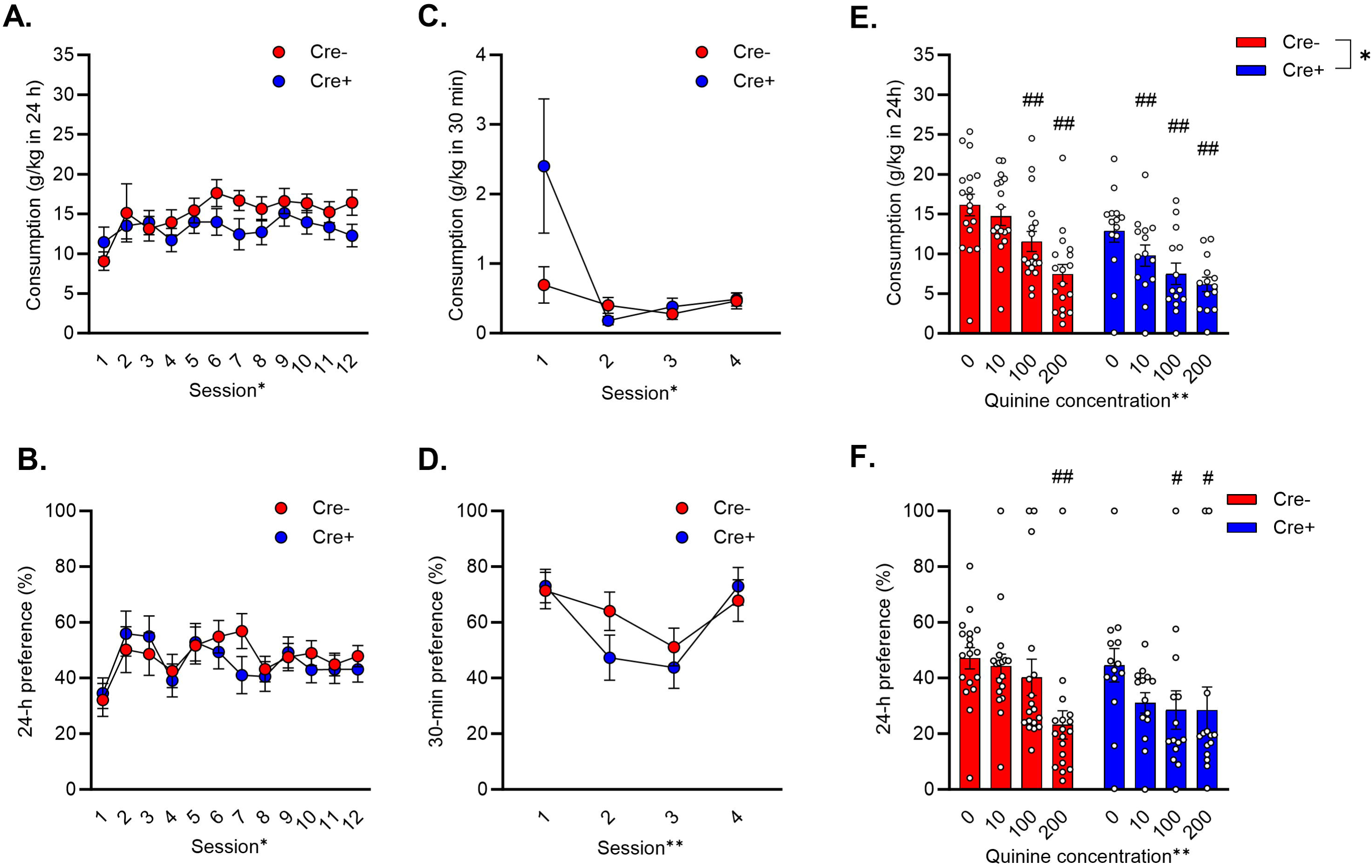
Mu-opioid receptor deletion on *Foxp2*-expressing neurons does not affect ethanol consumption or preference but does reduce aversion-resistant drinking. **A)** MOR deletion did not lead to differences in 24-h consumption or **B)** preference. **C)** Consumption of 20% EtOH during the first thirty minutes of intermittent access sessions was higher in Cre+ Foxp2-Cre/Oprm1^fl/fl^ mice on session 1. **D)** Preference did not differ between genotypes during the first thirty minutes of consumption. **E)** Compared to the 0-mg/L baseline sessions (= avg sessions 10-12), quinine reduced consumption of EtOH in Cre-control mice at concentrations of 100 and 200 mg/L and at all concentrations in Cre+ mice. Additionally, consumption was decreased in Cre+ mice vs. Cre-mice at all concentrations (main effect of genotype). **F)** Preference for EtOH vs. water was significantly reduced in Cre-mice at the 200 mg/L quinine concentration. In Cre+ mice, preference was reduced at the 100 and 200 mg/L quinine concentrations. * p < 0.05, ** p < 0.01 main effect (2-way ANOVA). # p < 0.05, ## p < 0.01 vs. 0 mg/L (Dunnett’s test).

Consumption and preference at 30 min were also measured for one session each week. Cre+ mice with *Foxp2*-MOR deletion were observed to consume more EtOH on the first 30-min session vs Cre-mice (**Figure 2C-D**). This was evidenced by a significant main effect of session [F_(1.057, 48.620)_ = 5.766, p < 0.05] and an interaction between session and genotype [F_(3, 138)_ = 3.131, p < 0.05] on consumption. There was no significant effect of genotype [F_(1, 46)_ = 2.053, p > 0.05]. For preference, RM two-way ANOVA revealed a significant main effect of session [F_(2.806, 129.1)_ = 6.100, p < 0.01], but no significant main effect of genotype [F_(1, 46)_ = 0.6369, p > 0.05] or session x genotype interaction [F_(3, 138)_ = 1.032, p > 0.05].

Aversion resistance was assessed by comparing consumption and preference on quinine sessions to the last three EtOH-only sessions (avg sessions 10-12). Foxp2-Cre/Oprm1^fl/fl^ Cre+ mice were more sensitive to quinine aversion than Cre-mice **(Figure 2E-F**). For consumption, RM two-way ANOVA found significant main effects of quinine concentration [F_(3, 90)_ = 43.150, p < 0.01] and genotype [F_(1, 30)_ = 4.588, p < 0.05]. The interaction of quinine concentration and genotype was not significant [F_(3, 90)_ = 2.314, p > 0.05]. A Dunnett’s multiple comparisons test between consumption at 0 mg/L and each quinine concentration was used to assess aversion resistance. There were significant decreases at 100 mg/L (p < 0.001) and 200 mg/L (p < 0.001) for Cre-animals. Cre+ mice significantly decreased consumption at 10 mg/L (p < 0.05), 100 mg/L (p < 0.001), and 200 mg/L (p < 0.001) (**Figure 2E**). When assessing EtOH preference, there was a significant main effect of quinine concentration [F_(3, 90)_ = 7.158, p < 0.01] but no effect of genotype [F_(1, 30)_ = 0.8399, p > 0.05] or interaction between the two [F_(3, 90)_ = 1.946, p > 0.05]. A Dunnett’s multiple comparisons test vs. baseline preference revealed significant decreases at 200 mg/L (p < 0.001) for Cre-animals and at 100 mg/L (p < 0.05) and 200 mg/L (p < 0.05) for Cre+ animals (**Figure 2F**). These results demonstrate that Foxp2-MOR deletion yields mice that are more sensitive to the suppressive effects of quinine on alcohol consumption and preference.

Water consumption on the intervening sessions (T/Th) was also evaluated and no effects of genotype were observed (**Figure 3A**). There was a significant main effect of session [F_(1.954, 89.890)_ = 5.789, p < 0.01] but no significant main effect of genotype [F_(1, 46)_ = 3.151, p > 0.05] nor an interaction between the two [F_(9, 414)_ = 1.421, p > 0.05].

**Figure 3.**
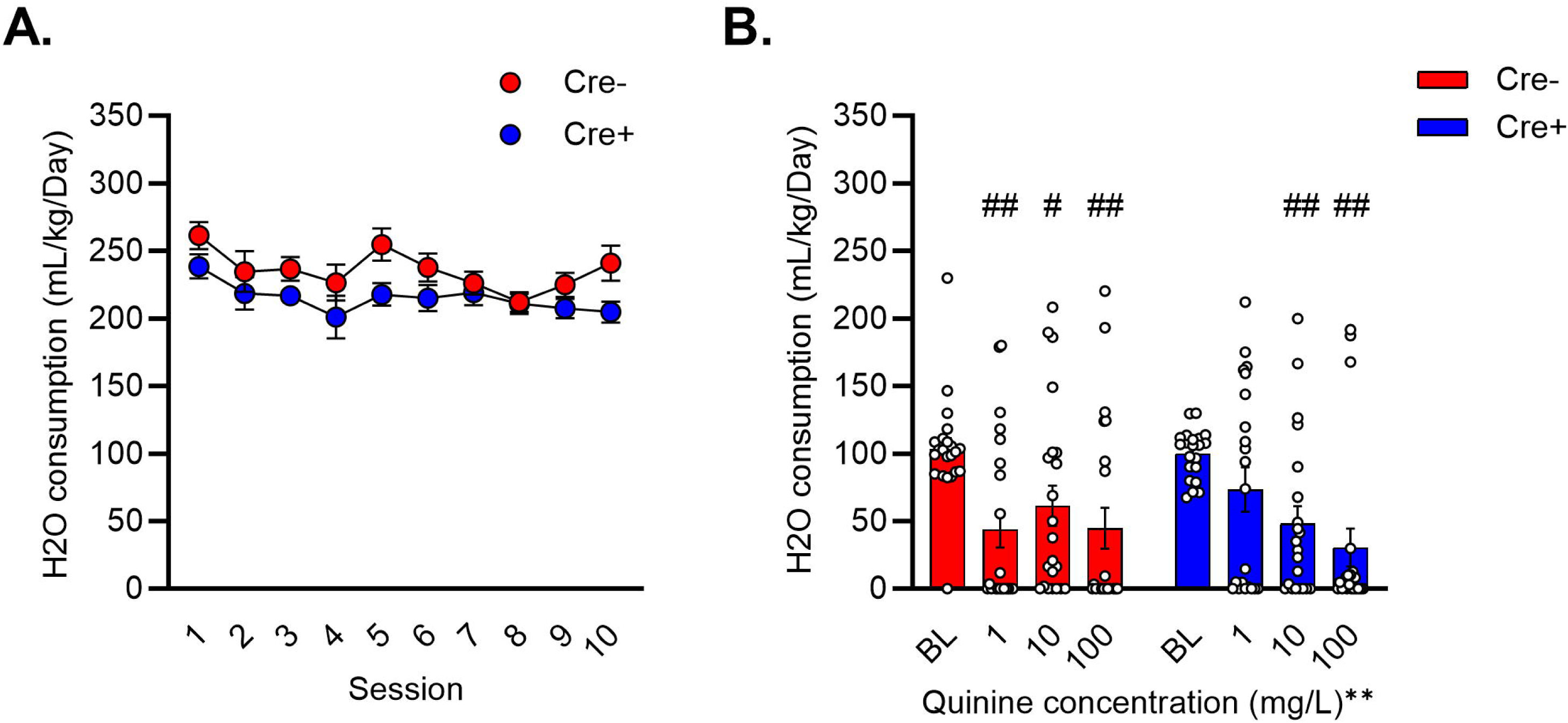
Mu-opioid receptor deletion on *Foxp2*-expressing neurons alters sensitivity to quinine in water. **A)** Water consumption on intervening days of the intermittent access paradigm did not differ between Cre- and Cre+ Foxp2-Cre/Oprm1^fl/fl^ mice. **B)** Cre-controls decreased water consumption at all quinine concentrations (1, 10, and 100 mg/L) while Cre+ mice with Foxp2-MOR deletion only reduced consumption at 10 and 100 mg/L. ** p < 0.01 main effect (2-way ANOVA). # p < 0.05, ## p < 0.01 vs. 0 mg/L (Dunnett’s test).

Quinine sensitivity was also assessed one week following the final week of EtOH consumption. Differences in quinine sensitivity were observed, with Cre+ mice appearing to be less sensitive to quinine than Cre-controls (**Figure 3B**). A mixed effects analysis found a significant main effect of quinine concentration [F_(3, 125)_ = 11.810, p < 0.0001] but not genotype [F_(1, 42)_ = 0.010, p > 0.05]. The interaction between concentration and genotype was not significant [F_(3, 125)_ = 1.649, p > 0.05]. A Dunnett’s multiple comparisons test used to assess changes from baseline revealed significant decreases at 1 mg/L (p < 0.001), 10 mg/L (p < 0.05), and 100 mg/L (p < 0.001) for Cre-animals, while Cre+ mice significantly decreased consumption at 10 mg/L (p < 0.01) and 100 mg/L (p < 0.001) only.

### 3.2. Operant responding for sucrose

All Foxp2-Cre/Oprm1^fl/fl^ mice learned to respond for sucrose at the active nose-poke hole with no difference in response rates between Cre- and Cre+ mice at any concentration (**Figure 4A**). A three-way RM ANOVA found significant main effects of sucrose concentration [F_(2, 120)_ = 26.58, p < 0.001] and response [F_(1, 60)_ = 425.1, p < 0.001] along with a sucrose concentration x response interaction [F_(2, 120)_ = 34.72, p < 0.001]. The main effect of genotype was not significant [F_(1, 60)_ = 0.05783, p > 0.05] and there were no significant interaction effects for sucrose concentration x genotype [F_(2, 120)_ = 0.3984, p > 0.05], response x genotype [F_(1, 60)_ = 1.834, p > 0.05] or sucrose concentration x response x genotype [F_(2, 120)_ = 1.601, p > 0.05]. A follow-up two-way RM ANOVA examining active responses across concentrations revealed a significant effect of sucrose concentration [F_(1.851, 55.53)_ = 34.77, p < 0.001] but not genotype [F_(1, 30)_ = 0.7063, p > 0.05]. There was also no significant interaction between sucrose concentration and genotype [F_(2, 60)_ = 0.9462, p > 0.05]. *Post-hoc* Holm-Sidak’s multiple comparisons tests revealed significant decreases in active responses at the 2% concentration (vs. 5% and 10%, p < 0.001 for both).

**Figure 4.**
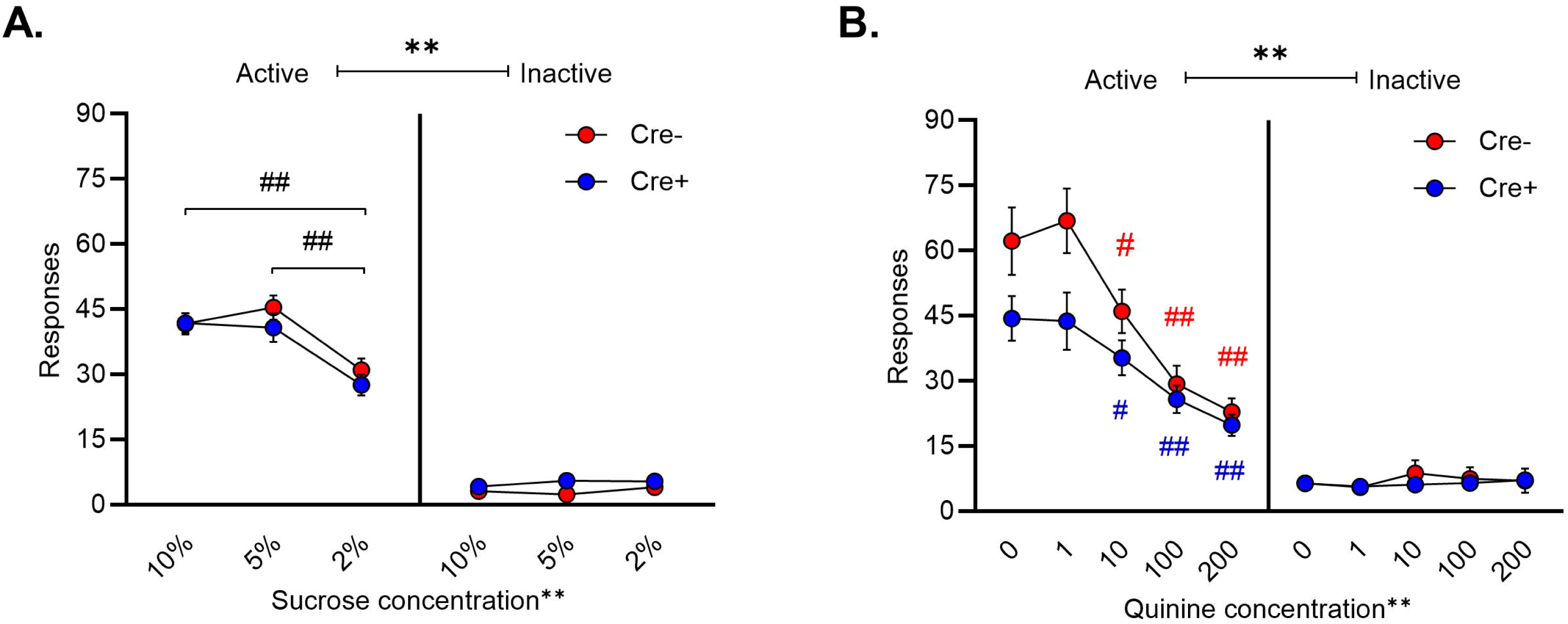
Operant responding for sucrose in mice with mu-opioid receptor deletion on *Foxp2*-expressing neurons. **A)** Foxp2-Cre/Oprm1^fl/fl^ mice were trained to respond for sucrose on the active nose-poke hole but responding was not affected by genotype. For both genotypes, active responses were lower for 2% vs. 10% and 5% sucrose. **B)** Cre- and Cre+ mice significantly decreased active responding for 2% sucrose at quinine concentrations of 10, 100, and 200 mg/L. For active responses, there was a significant quinine concentration x genotype interaction. # p < 0.05, ## p < 0.01 vs. 0 mg/L (Dunnett’s test).** p < 0.01 main effect (3-way ANOVA). ## p < 0.01 (Holm-Sidak test).

When analyzing sucrose responding in the presence of quinine, influences of genotype were uncovered (**Figure 4B**). A three-way RM ANOVA revealed significant main effects of quinine concentration [F_(2.978, 172.7)_ = 36.36, p < 0.001] and response [F_(1, 58)_ = 106.4, p < 0.001]. There were also several significant interaction effects, including quinine concentration x response [F_(4, 232)_ = 41.67, p < 0.001], quinine concentration x genotype [F_(4, 232)_ = 3.154, p < 0.05], and quinine concentration x response x genotype [F_(4, 232)_ = 3.591, p < 0.01]. However, there was no significant main effect of genotype [F_(1, 58)_ = 3.702, p > 0.05], nor an interaction effect between responses and genotype [F_(1, 58)_ = 2.958, p > 0.05]. A follow-up two-way RM ANOVA examining active responses revealed a significant quinine concentration x genotype interaction [F_(4, 116)_ = 4.048, p < 0.01] along with a significant main effect of quinine concentration [F_(2.548, 73.89)_ = 47.59, p < 0.001]. There was no significant effect of genotype [F_(1, 29)_ = 3.574, p > 0.05] in this analysis. To assess aversion resistance, Dunnett’s multiple comparisons tests comparing active responses at each quinine concentration to baseline (= 0 mg/L) were used. These revealed significant decreases at 10 mg/L (p < 0.05), 100 mg/L (p< 0.001), and 200 mg/L (p < 0.001) for both Cre- and Cre+ animals, suggesting that the genotypes were equally sensitive to quinine-punished sucrose seeking. Two-way RM ANOVA examining inactive responses revealed no significant effects.

To better understand the significant interactions between genotype and quinine concentration, we compared responses on the 0 mg/L and 1 mg/L sessions using unpaired t-tests between the genotypes.

These sessions were notably different from training since the response requirement was increased to FR3 to promote responding. These tests revealed a significant difference in responses for sucrose with 1 mg/L quinine (p = 0.027) while the difference at the 0 mg/L concentration approached the threshold for significance (p = 0.062). Together, the analyses suggest that Foxp2-MOR deletion impairs responding for sucrose when an FR3, but not FR1, response requirement is used while the negative influences of quinine on sucrose seeking are not impacted.

### 3.3. Conditioned place aversion

Morphine withdrawal-induced conditioned place aversion was observed in Foxp2-Cre/Oprm1^fl/fl^ mice and did not differ between the genotypes (**Figure 5A**). Analysis of time spent on the withdrawal-paired side revealed that place aversion was successful in both genotypes, as evidenced by a main effect of session [F_(1, 12)_= 19.460, p < 0.001] but no effect of genotype [F_(1, 12)_ = 1.992, p > 0.05] or interaction between genotype and session [F_(1, 12)_ = 1.344, p > 0.05].

**Figure 5.**
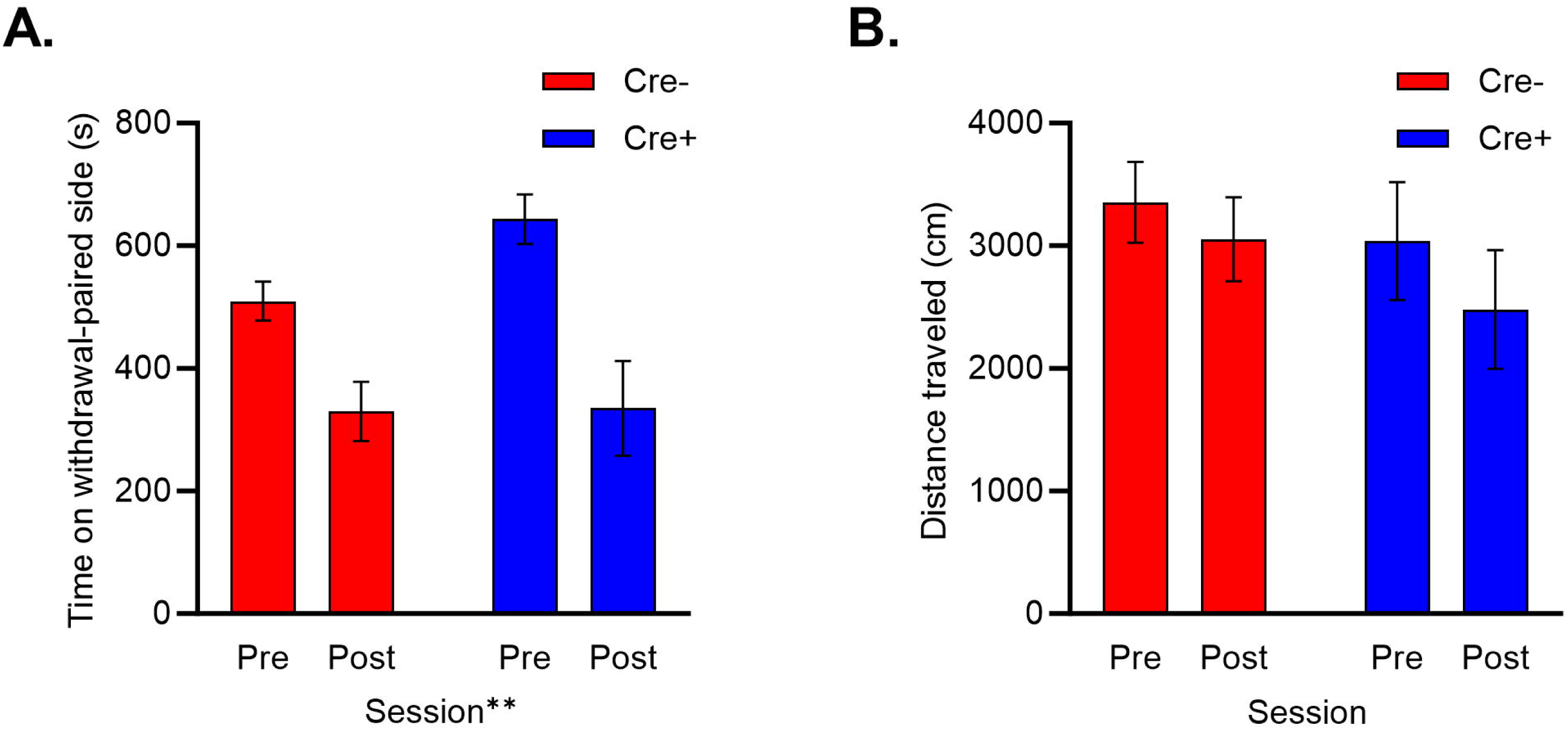
Mu-opioid receptor deletion on *Foxp2*-expressing neurons does not impact aversion to morphine withdrawal. **A)** Mice decreased time spent on the withdrawal-paired side during the post-vs. pre-conditioning test. The magnitude of aversion was not influenced by genotype. ** p < 0.01 main effect (2-way ANOVA). **B)** Distance traveled did not differ between sessions or genotypes.

When analyzing distance traveled during the test sessions, a two-way ANOVA revealed that neither genotype [F_(1, 12)_ = 0.671, p > 0.05] or session [F_(1, 12)_ = 3.607, p > 0.05] had a significant effect on locomotion (**Figure 5B**). There was also no interaction between genotype and session for this measure [F_(1, 12)_= 0.325, p > 0.05].

### 3.4. Morphine-induced locomotion

Reduced locomotion at baseline was observed in Cre+ (vs. Cre-) Foxp2-Cre/Oprm1^fl/fl^ mice. A two-way RM ANOVA on session 1 found no significant effects of genotype [F_(1, 21)_ = 2.380, p > 0.05] (**Figure 6A**). There was, however, a significant main effect of time [F_(2.218, 46.570)_ = 60.850 p < 0.001] and a time x genotype interaction [F_(5, 105)_ = 5.193, p < 0.001]. Following morphine injection, locomotor activity increased and still differed between Cre- and Cre+ mice. A two-way RM ANOVA on session 2 found no significant effects of genotype [F_(1, 21)_ = 3.068, p > 0.05] (**Figure 6B**). There was a significant main effect of time [F_(1.500, 31.510)_ = 14.660, p < 0.001], and a time x genotype interaction [F_(11, 231)_ = 2.724, p < 0.01].

**Figure 6.**
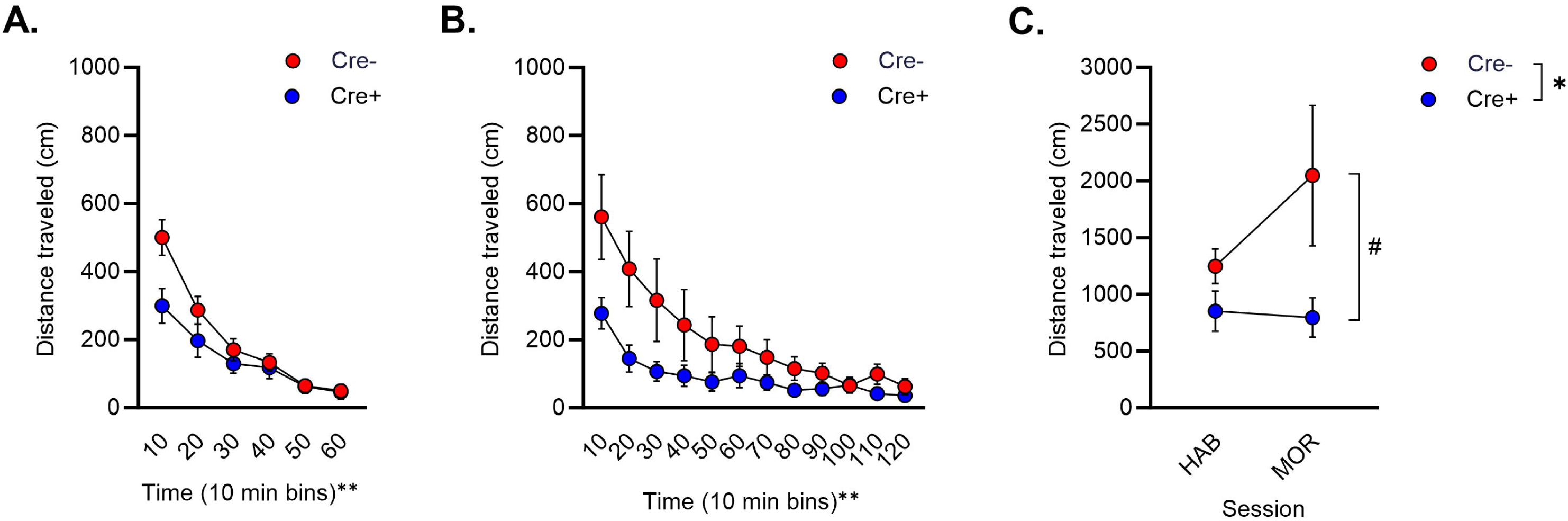
Locomotion is decreased by *Foxp2* mu-opioid receptor deletion. **A)** Locomotor activity was decreased in Cre+ Foxp2-Cre/Oprm1^fl/fl^ mice in a time-dependent manner (significant time x genotype interaction). **B)** Following a 20-mg/kg injection of morphine, locomotor activity was lower in Cre+ mice with MOR deletion in a time-dependent manner (significant time x genotype interaction). **C)** Morphine-induced increases in locomotion were greater in Cre-vs. Cre+ mice. Total distance in the first 60 min of the morphine challenge session is shown. ** p < 0.01 main effect (2-way ANOVA). # p < 0.05 (Holm-Sidak test).

To determine if morphine caused an unconditioned increase in locomotion in either genotype, distance traveled during the first 60 min of each session was totaled and compared using a two-way ANOVA. This analysis revealed a main effect of genotype [F_(1, 20)_ = 5.715, p < 0.05]. The main effect of session [F_(1, 20)_ = 1.199, p > 0.05] and session x genotype interaction [F_(1, 20)_ = 1.586, p > 0.05] were not significant. *Post hoc* Holm-Sidak’s multiple comparison tests found that locomotion was significantly greater in Cre-vs. Cre+ mice on session 2 (p < 0.05) only.

## 4. Discussion

Together, the results of these studies in mice with genetic deletion of *Oprm1* in *Foxp2*-expressing neurons suggest that this neuronal subpopulation contributes to the tendency to seek out or consume alcohol despite the presence of a concurrent aversive stimulus. Cre+ mice with *Foxp2*-MOR deletion were more sensitive to quinine punishment than their Cre-littermates when consuming alcohol in an intermittent access task. This pattern of results cannot be explained by differences in quinine sensitivity between Cre- and Cre+ mice, as the latter were actually less sensitive to quinine in water and decreased responding to the same quinine concentrations when it was presented in sucrose. We also found that Cre+ mice with MOR deletion tended to have lower active responses for sucrose than Cre-mice when an FR3 schedule was used. This result may reflect reduced motivation or willingness to exert effort in mice with *Foxp2*-MOR deletion. However, considering that EtOH drinking and responding for sucrose on an FR1 schedule remained intact, *Foxp2*-MORs do not appear to be necessary for simple responding for reward.

A number of prior studies have implicated the MOR in alcohol consumption through genetic models in which *Oprm1* is deleted non-specifically throughout the nervous system. For example, in an IA task in which consumption was measured twice per week, mice lacking MOR drank significantly less 10% EtOH compared to wild-type controls (Becker et al., 2002). Similarly, female heterozygous and homozygous MOR-knockout mice consumed significantly less EtOH than wild-type female mice when the EtOH concentration was below 15%, with no differences in male mice (Hall et al., 2001). MOR deletion targeted to GABAergic forebrain neurons, which includes many *Foxp2* neurons, also decreased consumption in mice consuming 20% EtOH in an IA task (Ben Hamida et al., 2019). The results of the current studies suggest that selective deletion of MOR on *Foxp2*-expressing neurons is not sufficient to replicate these effects, although it is important to consider that the impact of genotype could emerge at lower EtOH concentrations. It should also be noted that a main effect of genotype was observed for consumption during the quinine sessions, with Cre+ mice consuming less than Cre-littermates during these final six sessions. Further, although there was no statistically significant interaction between genotype and session during the IA experiment, consumption in Cre-mice increased by approximately 7 g/kg between sessions 1 and 12 while the increase in Cre+ mice was < 1 g/kg. Considering these trends, we speculate that MOR expression in the *Foxp2*-negative MSNs was sufficient to maintain consumption near control levels for the majority of the experiment but that *Foxp2* neurons may be important for escalation of alcohol consumption.

Despite the robust literature implicating the endogenous opioid system in alcohol-drinking behaviors, there have been only a few investigations into whether or how this system contributes to aversion-resistant drinking. Systemic MOR antagonism has been shown to reduce alcohol seeking in rats that are resistant to a footshock punishment (Giuliano et al., 2018), and MOR internalization and recycling have been linked to the development of compulsive morphine-seeking (Berger and Whistler, 2011). Our group has also recently found that deletion of *Oprm1* in cholinergic neurons, which includes striatal cholinergic interneurons that regulate dopamine release onto MSNs, reduces sensitivity to the suppressive effects of quinine on alcohol consumption (Beane et al., 2024). The results detailed here further suggest a role for MOR in aversion-resistant alcohol drinking and also highlight the potential importance of *Foxp2*-expressing neurons to this behavior.

As noted, *Foxp2* neurons are found throughout the forebrain, including in the amygdala, striatum, and cortex. Considering the role of the ventral striatum in regulating risk and punishment, the effects of *Foxp2*-MOR deletion on aversion-resistant drinking may rely on this nucleus (Piantadosi et al., 2021). For instance, glutamatergic inputs from the mPFC and insula to the NAc core and shell, but not the basolateral amygdala, are key regulators of this behavior (Halladay et al., 2020; Seif et al., 2013). Non-specific inhibition of the NAc core also reduced quinine-resistant drinking (Sneddon et al., 2021). However, because aversion resistance was not impacted by targeted inhibition of D1-expressing or D2-expressing neurons in the NAc core, an alternative subpopulation of striatal neurons must be regulating the behavior. Based on our recent work, *Foxp2*-expressing MSNs or cholinergic interneurons (Beane et al., 2024) may serve this function. A role for cortical *Foxp2* neurons also cannot be excluded, as this population has been implicated in reversal learning, another form of behavioral flexibility (Co et al., 2020).

In addition to observing differences in alcohol drinking behaviors, Cre+ Foxp2-Cre/Oprm1^fl/fl^ mice exhibited a number of behavioral differences that mimicked those observed in animals with non-specific MOR deletions. First, there was a trend for Cre+ mice to respond less for sucrose (although only when the response requirement was increased from FR1 to FR3). Reduced responding for a sucrose reward, but not a food pellet, has previously been observed in mice with systemic MOR deletion (Roberts et al. 2000). Reduced locomotion, both at baseline and in response to a morphine challenge, has also been observed previously in MOR knockout mice (Hall et al., 2003). Replication of this result following *Foxp2*-MOR deletion is not surprising as approximately 70% of striatal *Foxp2* neurons express D1 receptors, which can stimulate locomotion (Desai et al., 2005; Gong et al., 1999; Tran et al., 2005) and are critical for the locomotor activating effects of opioids (Becker et al., 2001; Urs et al., 2011). Mutations in *Foxp2* also cause severe motor impairments in mice (French and Fisher, 2014; Fujita et al., 2008).

Importantly, these changes in locomotor activity did not impact the ability of Cre+ Foxp2-Cre/Oprm1^fl/fl^ mice to engage in the alcohol-drinking or operant response tasks and differences in locomotion were not observed in the place conditioning experiment. Considering that the differences in activity levels occurred in the first 10-20 minutes of the sessions, they may be specific to novelty-induced locomotion and would not confound the other effects reported here. Finally, mice with *Foxp2*-MOR deletion were able to learn to respond for reward and to form associations with morphine withdrawal. These results align with prior conclusions that non-specific MOR deletion produces deficits in motivation but not learning processes (Kas et al., 2004; Lubbers et al., 2007).

A limitation of this work is that *Foxp2* is expressed in many types of cells throughout the nervous system. Thus, we cannot conclude whether the effects observed here are due to *Foxp2*-expressing MSNs, cortical neurons, amygdala ITC cells, or another population of neurons. The influences of these subpopulations are further confounded by the fact that they are interconnected. For example, around 23% of neurons in the apical ITC (apITC) cluster project into the striatum, where apITC cells release GABA to inhibit striatal mechanisms that regulate amygdala-striatal behavioral responses (Asede et al., 2022, 2021). Cortico-striatal projection neurons are also known to express *Foxp2* (French and Fisher, 2014; Hisaoka et al., 2010). Future studies on the role of MORs in aversion resistance should strive for increased region and cell-type specificity to better understand the role of these receptors in reward-seeking and consuming behaviors despite adverse consequences.

## Conclusions

Together, these investigations demonstrate that targeted genetic deletion of MOR on *Foxp2* neurons does not affect alcohol consumption or preference but may prevent escalation of consumption with time and exposure. Mice with *Foxp2*-MOR deletion also fail to develop aversion-resistant alcohol drinking, suggesting that this subpopulation of neurons may be important to the development of the maladaptive, compulsive behaviors characteristic of addiction.

## Acknowledgments

The authors are grateful to Kiara Ream, Ezra Eccles, and Caroline Scribner for assistance with behavioral experiments.

## Author contributions

Harrison Carvour: Conceptualization, Formal Analysis, Investigation, Writing - Original Draft, Visualization. Charlotte Roemer: Investigation, Writing - Original Draft. D’Erick Underwood: Investigation, Writing - Original Draft. Edith Padilla: Formal Analysis, Investigation, Writing - Original Draft. Oscar Sandoval: Investigation, Writing - Original Draft. Megan Robertson: Formal Analysis, Investigation, Writing - Original Draft. Mallory Miller: Formal Analysis, Investigation. Natella Parsadanyan: Formal Analysis, Investigation. Thomas Perry: Investigation, Visualization. Anna Radke: Conceptualization, Formal Analysis, Writing - Review and Editing, Visualization, Supervision, Funding Acquisition.

